# Bolstering fitness via opportunistic CO_2_ fixation: mixotroph dominance in modern groundwater

**DOI:** 10.1101/2021.01.26.428071

**Authors:** Martin Taubert, Will A. Overholt, Beatrix M. Heinze, Georgette Azemtsop Matanfack, Rola Houhou, Nico Jehmlich, Martin von Bergen, Petra Rösch, Jürgen Popp, Kirsten Küsel

## Abstract

The current understanding of organic carbon inputs into ecosystems lacking photosynthetic primary production is predicated on data and inferences derived almost entirely from metagenomic analyses. The elevated abundances of putative chemolithoautotrophs in groundwaters suggest that dark CO_2_ fixation is an integral component of subsurface trophic webs. To understand the impact of autotrophically-fixed carbon, the flux of CO_2_-derived carbon through various subpopulations of subsurface microbiota must first be resolved, both quantitatively and temporally. Here, we implement novel stable isotope cluster analysis to render a time-resolved and quantitative evaluation of ^13^CO-derived carbon flow through a groundwater microbiome stimulated with reduced sulfur compounds. We demonstrate that mixotrophs, not obligate chemolithoautotrophs, were the most abundant active organisms in groundwater microcosms. Species of *Hydrogenophaga*, *Polaromonas*, *Dechloromonas*, and other metabolically versatile mixotrophs drove the recycling of organic carbon and, when chance afforded, supplemented their carbon requirements via chemolithoautotrophy and uptake of available organic compounds. Mixotrophic activity facilitated the replacement of 43 and 80% of total microbial carbon stores with ^13^C in just 21 and 70 days, respectively. This opportunistic “utilize whatever pathways net the greatest advantage in fitness” strategy may explain the great abundances of mixotrophs in other oligotrophic habitats, like the upper ocean and boreal lakes.

---

From soils to deep-sea sediments, the vast majority of cells on Earth must find a way to thrive in environments devoid of photosynthesis^1^. To truly appreciate the global carbon cycle in all its grandeur, it is important to understand the extent to which various types of cells rely upon allochthonous or autochthonous carbon input. This dependence invokes selective pressures that favor heterotrophic or chemolithoautotrophic lifestyles and provides the foundation upon which trophic webs linking the entire subsurface biome are structured. Accurately gauging CO_2_ fixation rates and turnover in these habitats is remarkably challenging despite the invaluable utility afforded by metagenomics to shed light on the metabolic capabilities of thousands of the organisms present^2–5^.

Modern groundwater, i.e., water having ingressed into the subsurface within the past 50 years^6^, is a transitionary ecosystem that connects surface habitats dominated by recently photosynthetically fixed carbon with the subsurface, which is devoid of this carbon source entirely^7–9^. Here, inorganic electron donors like reduced nitrogen, iron, and sulfur fuel chemolithoautotrophic primary production^9–12^. Metagenomic-based studies have elucidated a diverse array of microorganisms bearing the metabolic potential for chemolithoautotrophy^4, 13–16^, accounting for 12 to 47% of the microbial population detected in groundwater^17–20^. Discoveries like these have cast doubt on paradigms portraying modern groundwater as being dominated by heterotrophic microbes fueled by organic material from the surface. We hypothesize that chemolithoautotrophic primary production dictates the rates by which carbon is cycled in the modern groundwater microbiome.

To validate this hypothesis, we implemented a novel approach - stable isotope cluster analysis (SIsCA), to render a time-resolved, quantitative assessment of CO_2_-derived carbon flow through the groundwater food web. By coupling stable isotope probing (SIP) with genome-resolved metaproteomics^21,22^, we leveraged the high sensitivity of Orbitrap mass spectrometry in SIP-metaproteomics to acquire exceedingly accurate quantitative data on ^13^C incorporation^23,24^. SIsCA then employs a dimensionality reduction approach to incorporate the molecule-specific temporal dynamics of the acquired isotopologue patterns resulting from ^13^C-SIP time-series experiments – ultimately discerning trophic interactions between individual members of the microbial community.

To examine the role of chemolithoautotrophy in the groundwater microbiome, we amended groundwater microcosms with ^13^CO. Thiosulfate was used as an electron donor, as it is regularly released into groundwater via rock weathering^25–28^. While organisms bearing the genetic potential to oxidize reduced sulfur compounds are widespread in groundwater, little is known about their lifestyles^15,16,29^. Under conditions favoring lithotrophic growth, we expected chemolithoautotrophy to be the primary source of organic carbon, and a unidirectional carbon flux from autotrophs to heterotrophs. By mapping the quantitative information derived from SIsCA to MAGs, we were able to characterize carbon utilization and trophic interactions between active autotrophs and heterotrophs in the groundwater microbiome over a period of 70 days. High-resolution monitoring of carbon cycling and taxon-specific activities demonstrated that metabolically versatile mixotrophs, not strict autotrophs, drove carbon flux in the groundwater, supplying up to 80% of the entire microbial carbon. Insights into these microbes’ lifestyles, as well as a discussion on how a metabolically flexible mixotrophic lifestyle is optimally fit to flourish in an oligotrophic ecosystem, ensues.

## Materials and Methods

### Groundwater sampling and microcosms setup

Groundwater was collected from Hainich Critical Zone Exploratory (CZE) well H41 (51.1150842N, 10.4479713E) in June 2018. Well H41 provides access to an aquifer assemblage at 48 m depth in a trochite limestone stratum. Sourced by a beech forest (*Fagus sylvatica*) recharge area, the oxic groundwater in this well maintains mean dissolved oxygen concentrations of 5.0 ± 1.5 mg L^−1^, < 0.1 mg L^−1^ ammonium, 1.9 ± 1.5 mg L^−1^ dissolved organic carbon, 70.8 ± 12.7 mg L^−1^ total inorganic carbon, and a pH of 7.2 ^27,30^. A total of 120 L of groundwater was sampled using a submersible pump (Grundfos MP1, Grundfos, Bjerringbro, Denmark). To collect biomass from the groundwater, 5-liter fractions were filtered through each of twenty 0.2-μm Supor filters (Pall Corporation, Port Washington, NY, USA). The natural background of inorganic carbon in the groundwater was then replaced with defined concentrations of ^12^C or ^13^C. Two 3-liter volumes of filtered groundwater were acidified to pH 4 in 5-liter bottles to eliminate any bicarbonate. Following that, ^12^C- or ^13^C-bicarbonate was dissolved in the groundwater to a final concentration of 400 mg L^−1^, corresponding to a near *in situ* concentration of 79 mg C L^−1^. The pH of groundwater samples was then adjusted to 7.2 by addition of ^12^C- or ^13^C-CO_2_.

Eighteen distinct microcosms were initiated for the ^13^C-SIP experiment. For each microcosm, one sample-laden 0.2-μm filter was placed into a 500-mL bottle containing 300 mL of treated groundwater (as described above). Nine microcosms were sourced with water containing ^12^C-bicarbonate and the other nine with water containing ^13^C-bicarbonate. Two additional microcosms were prepared, each by transferring one 0.2-μm filter into a 1-liter bottle containing 350 mL of untreated groundwater. One of these bottles was supplemented with 150 mL sterile D_2_O (final concentration 30%, v:v) and the other with 150 mL sterile milliQ H_2_O. Sodium thiosulfate and ammonium chloride were then was added to all microcosms to a final concentration of 2.5 mM and 15 μM, respectively. Finally, all microcosms were incubated with shaking (100 rpm) at 15 °C in the dark.

### Hydrochemical analyses

While incubating the 18 microcosms supplemented with ^12^C- or ^13^C-bicarbonate, concentrations of oxygen, thiosulfate, and sulfate were determined at regular intervals. Oxygen concentrations were determined using a contactless fiber-optic oxygen sensor (Fibox 4 trace with SP-PSt3-SA23-D5-YOP-US dots [PreSens Precision Sensing GmbH, Regensburg, Germany]). Measurements were collected from three ^12^C microcosms and three ^13^C microcosms every two days for the first three weeks, and once weekly thereafter. Thiosulfate concentrations were determined via colorimetric titration assays with iodine^31^. Samples from all microcosms were evaluated every four to seven days. For each measurement, 2 mg potassium iodide was mixed into 1 mL of sample, followed by the addition of 10 μL of zinc iodide-starch solution (4 g L^−1^ starch, 20 g L^−1^ zinc chloride and 2 g L^−1^ zinc iodide) and 10 μL of 17% (v:v) phosphoric acid. Titration was performed by adding 5 μL of 0.005 N iodine at a time until the solution turned faint blue. Thiosulfate concentrations (*c_thiosulfate_* in mg L^−1^) were then calculated according to equation (1), where *V_iodine_* is the volume of iodine solution added and *V_sample_* is the sample volume:

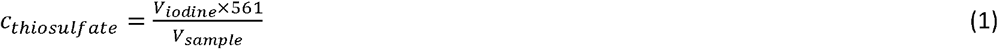

Sulfate concentrations were determined via a turbidimetric assay^32^ from all microcosms every four to seven days. For each measurement, 1 mL of either microcosm sample, standard (50 μM to 1000 μM potassium sulfate) or blank (dH_2_O) was mixed with 0.4 mL 0.5 M HCl and 0.2 mL BaCl_2_-gelatin reagent (0.5 g gelatin and 8 g BaCl_2_ in 200 mL dH_2_O). Following 1 h incubation in the dark, absorbances were measured at 420 nm in a DR3900 spectrophotometer (HACH, Düsseldorf, Germany).

### Detection of cellular activity by Raman microspectroscopy

Microcosms supplemented with D_2_O or H_2_O were sampled regularly during the first seven weeks of incubation to quantify the incorporation of deuterium into the biomolecules of active cells (i.e., carbon-deuterium [C-D] bonds) via single-cell Raman microspectroscopy analysis. In preparation for Raman microspectroscopy, 1 mL of sample was pre-filtered through a 5-μm filter, and then the cells contained in the filtrate were washed three times with ddH_2_O via centrifugation (10,000g, 2 min). Pellets were then resuspended in 50 μL ddH_2_O, and 10 μL of the final suspension was placed on nickel foil (Raman substrate) and allowed to air dry at RT. Microbial cells were located via dark field microscopy, and measurements were collected using a Raman microscope (BioParticleExplorer 0.5, rap.ID Particle Systems GmbH) with an excitation wavelength of 532 nm (solid-state frequency-doubled Nd:YAG module [Cobolt Samba 25 mW]; laser power = 13 mW at sample). The laser was focused with an x100 objective (Olympus MPLFLN 100xBD) across a lateral spot of < 1 μm. Backscattered light (180°) was diffracted using a single-stage monochromator (Horiba Jobin Yvon HE 532) with a 920 line mm^−1^ grating. Spectra were then registered with a thermoelectrically cooled CCD camera (Andor DV401-BV), resulting in a resolution of ~ 8 cm^−1^. A 5 s integration period was applied per Raman spectrum (−57 to 3203 cm^−1^).

### Processing and analysis of Raman data

Processing and statistical analysis of raw Raman data were achieved with GNU R software^33^. Cosmic spikes were removed from the spectra^34^. A wavenumber calibration was then applied using 4-acetamidophenol standard spectra^35^, while an intensity calibration was performed using the SRM2242 standard^36,37^. The contribution of fluorescence was removed from spectra using the asymmetric least-squares baseline correction method^38^. Finally, spectra were vector-normalized and subjected to dimensionality reduction via principal component analysis (PCA). Five principal components were used to build a linear discriminant analysis classification model, which was applied to differentiate between deuterium-labeled and unlabeled bacterial cells. Deuterium uptake was expressed as the C-D ratio, i.e., A(C-D) / [A(C-D) + A(C-H)], which was calculated by integrating the areas of the C-H (2800 - 3100 cm^−1^) and C-D (2040 - 2300 cm^−1^) stretching vibration bands. Monitoring deuterium incorporation into microbial cells helped gauge metabolic activity, as well as determine optimal time points to sample microcosms.

### Sampling and biomolecule extraction

After 21, 43, and 70 days of incubation, biomass was recovered from microcosms by filtering aqueous phases through 0.2-μm Supor filters (Pall Corporation). Filters used for pre-incubation biomass enrichment were combined with the filters used to remove the aqueous phases. A combined DNA and protein extraction was performed using a phenol/chloroform/isoamylalcohol-based protocol, as previously described^39^. Details regarding 16S rRNA gene amplicon sequencing and quantitative SIP of DNA are provided in Supp. Info.

### Metagenomic analysis

Metagenomic sequencing was performed on DNA samples selected from four ^12^C microcosms: one replicate each following 21 and 43 days of incubation, and two replicates following 70 days of incubation. Samples were selected with the aim of covering greatest taxonomic diversity, as per the results of PCA of 16S rRNA gene amplicon sequencing data. DNA fragment sizing, quantitation, integrity, and purity were determined using an Agilent 2100 Bioanalyzer (Santa Clara, CA, USA). Library preparation was achieved with a NEBNext Ultra II DNA Lib Prep Kit (New England Biolabs, Ipswich, MA, USA) in accordance with protocols provided by the manufacturer. Multiplexed sequencing in one flow cell of an Illumina NextSeq 500 system (300 cycles) ensued to generate 150-bp paired-end reads.

Raw sequencing data was quality filtered using BBDuk^40^ and subjected to assembly with metaSPAdes v3.13.0^41^. Applying only contigs greater than 1,000 bp in length, three different algorithms facilitated genomic binning: MaxBin 2.0 v2.2.7^42^, MetaBAT 2 v2.12.1^43^, and BinSanity v0.2.7^44^. Bin refinement was accomplished using the MetaWRAP pipeline v1.1.3^45^. Only bins that were both more than 50% complete and contained less than 10% contamination were considered. Bins were classified with GTDB-Tk v0.3.2^46^, and completeness parameters were appraised with CheckM v1.0.12^47^. Bins from different samples were dereplicated using FastANI v1.0^48^. The Prokka pipeline v1.13.3^49^ was used to assign functional annotations to gene sequences and to translate into amino acid sequences for metaproteomics analysis. Metagenomic bins of particular interest (per metaproteomics analysis) were manually refined with Anvi’o v6.1^50^, rendering the completed MAGs. Normalized coverage values for all MAGs were calculated by dividing raw coverage values by the relative abundance of *rpoB* genes in each metagenome. Gene abundances of *rpoB* were determined using ROCker^51^. Table S1 provides an overview of the curated MAGs referred to in the study.

### Metaproteomics analysis

Proteins extracted from microcosms were first subjected to SDS polyacrylamide gel electrophoresis, followed by in-gel tryptic cleavage as previously described^39^. After reconstitution in 0.1% formic acid (v:v), LC-MS/MS analysis was performed in LC chip coupling mode on a Q Exactive HF instrument (Thermo Fisher Scientific, Waltham, MA, USA) equipped with a TriVersa NanoMate source (Advion Ltd., Ithaca, NY, USA). Raw data files were analyzed using the Sequest HT search algorithm in Proteome Discoverer (v1.4.1.14, Thermo Fisher Scientific, Waltham, MA, USA). Amino acid sequences derived from the translation of genes present in metagenomes were compiled into a reference database to facilitate protein identification. The following parameters were applied: enzyme specificity was set to trypsin, two missed cleavages were allowed, oxidation (methionine) and carbamidomethylation (cysteine) were selected as modifications, and peptide ion and Da MS/MS tolerances were set to 5 ppm and 0.05, respectively. Peptides were considered identified upon scoring a q-value < 1% based on a decoy database and obtaining a peptide rank of 1. Functional classification of peptides was achieved in accordance with gene annotations generated by Prokka^49^, and taxonomic classification was based on the dereplicated and refined MAGs described above.

### Stable Isotope Cluster Analysis

Peptide identifications from ^12^C microcosms samples were mapped to mass spectra of corresponding ^13^C-labeled samples, and the incorporation of ^13^C was quantified by comparing expected peptide masses, chromatographic retention times, and MS/MS fragmentation patterns. Molecular masses of peptides were calculated based on amino acid sequences, isotopologue patterns of labeled peptides were extracted manually from mass spectral data using the Xcalibur Qual Browser (v3.0.63, Thermo Fisher Scientific, Waltham, MA, USA), and ^13^C incorporation was calculated as previously described^24^.

The conventional approach of calculating the most probable ^13^C relative isotope abundance (RIA) of a peptide does not take into account the information contained in isotopologue patterns, which provide detailed information about the carbon utilization of an organism. To include this information in the analysis, we developed Stable Isotope Cluster Analysis (SIsCA). Stable Isotope Cluster Analysis (SIsCA) was performed using R^33^, with scripts being available on github (https://github.com/m-taubert/SIsCA). Measured isotopologue patterns were compared to 21 predicted isotopologue patterns varying in ^13^C RIA (5% intervals from 0 to 100% ^13^C RIA), and coefficients of determination (R^2^) were calculated for each comparison. With this approach, all information from isotopologue patterns is retained, while still data from different peptides is comparable, time series can be integrated, and the dataset can easily be used for downstream statistical analysis. To differentiate microbial lifestyles, R^2^ values were averaged from samples obtained from replicate microcosms and peptides assigned to the same MAG. Resulting datasets of 21 R^2^ values per time point per MAG were compared via PCA with the vegan software package^52^, and clusters of MAGs were defined manually and validated by testing for overlapping confidence intervals.

Generation times of individual taxa were calculated by comparing the relative intensity of unlabeled and labeled peptide signals in mass spectrometric data, as previously described^23^. The number of doublings, *n*, was calculated according to equation (2) where 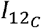 and 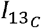 are the signal intensities of the unlabeled peptide and labeled peptide, respectively:

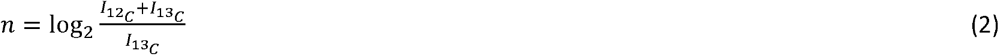

If the mass spectrometric signals of unlabeled and labeled and peptides overlapped, the monoisotopic peak was used to determine the total abundance of unlabeled peptide based on the natural distribution of heavy isotopes, as previously described^24^. Generation time, *t_d_*, was calculated with equation (3), where Δ*t* is incubation time:

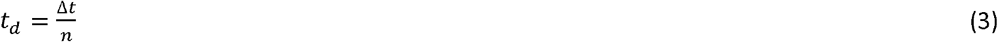

## Results

### Sulfur oxidation by active groundwater microbes

Groundwater microbiota responded immediately to the addition of thiosulfate, yielding increasing rates of sulfur oxidation. During the first three weeks of incubation, thiosulfate and oxygen consumption rates remained relatively low (1.7 ± 1.9 and 5.5 ± 2.0 μmol d^−1^ [mean ± SD], respectively; Fig. S1). Raman microspectroscopic analyses suggested that > 95% of cells were active within the first 12 days of incubation. A distinct C-D band was observed at wavelength positions between 2,100 and 2,300 cm^−1^ in the single-cell Raman spectra of the microcosm amended with D_2_O (Fig. 1, Fig. S2), which demonstrated new biomolecules were being synthesized by incorporating deuterium from D_2_O into carbon-deuterium bonds. The relative intensity of the C-D band increased from 18.3% after 12 days to 25.7% after 47 days of incubation (median values; *p* < 2.2 × 10^−16^, *t* = − 14.038, *df* = 141.68, two-sided Welch’s *t*-test), indicative of continued microbial proliferation and cross-feeding on deuterium-labeled organic carbon.

**Figure 1:**
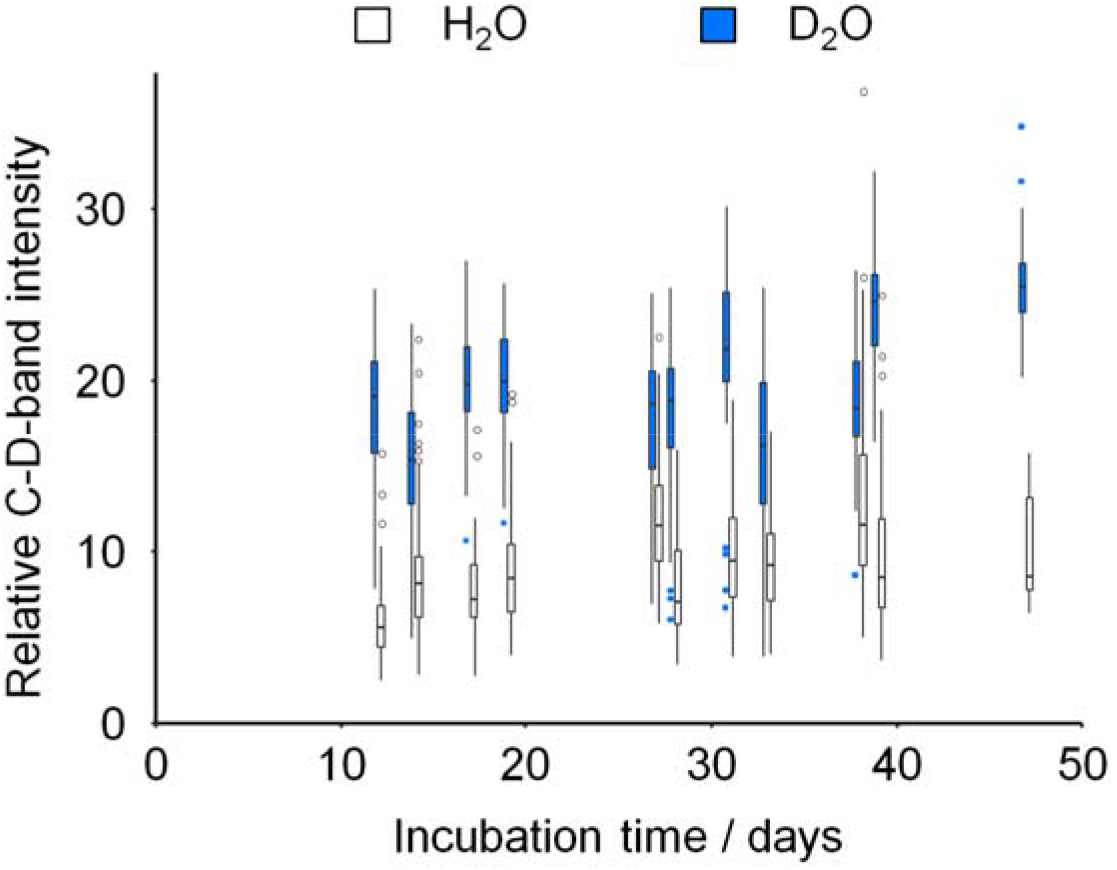
Quantification of deuterium incorporation by single-cell Raman microspectroscopy. Boxplots depict the relative intensity of Raman C-D bands, determined by A(C-D) / [A(C-D) + A(C-H)], from single-cell Raman spectra. Spectra were obtained from groundwater microcosms with 30% D_2_O (shaded) or H_2_O (empty) at various time points. Boxes show median, and first and third quartile; whiskers denote 5^th^ and 95^th^ percentile. Outliers are depicted as dots. A minimum of 147 spectra were obtained at each time point.

After 70 days of incubation, consumption rates of thiosulfate (7.2 ± 2.0 μmol d^−1^) and oxygen (12.8 ± 3.2 μmol d^−1^) had increased significantly (*p* = 6.48 × 10^−4^, *t* = 5.4332, *df* = 7.8999 [thiosulfate] and *p* = 1.27 × 10^−3^, *t* = 4.7692, *df* = 8.3019 [oxygen], two-sided Welch’s t-test, Fig. S1). Sulfate was produced at a consistent rate ranging between 8.1 and 9.6 μmol d^−1^ (no significant changes) throughout the duration of the experiment. Recorded stoichiometry for oxygen:thiosulfate:sulfate was roughly 2.8 : 1 : 2.6 over the course of incubation, very near the theoretical ratio of 2 : 1 : 2 for oxygen-dependent thiosulfate oxidation.

### Organism-specific ^13^C incorporation reveals distinct lifestyles

To address the carbon utilization schemes of key microbes, we conducted genome-resolved SIP-metaproteomic analyses after 21, 43, and 70 days of incubation. SIsCA then clustered the 31 most abundant MAGs into five distinct groupings, based on carbon utilization (Fig. 2, Fig. S3, Dataset S1). Organisms represented by MAGs in cluster I were related to *Thiobacillus* (*Burkholderiales*) and exhibited a stable ^13^C RIA of 95% over throughout the 70 day experiment (Fig. 2). Such a high (>90%) ^13^C RIA indicated exclusive CO_2_ fixation. However, these strict autotrophs accounted for only 11% of the total number of MAGs across the five clusters and 3.2 ± 3.1% (mean ± sd) of the total biodiversity enveloped by the community, based on normalized coverages of the metagenomics dataset and 16S rRNA gene profiles, respectively (Fig. S4, Fig. S5, Supp. Info.). By comparing the signal intensities of ^12^C- and ^13^C-enriched peptides, the generation time of these autotrophs was determined to be less than 2 days (Fig. 3), highlighting the rapid production of new ^13^C-labeled biomass from ^13^CO_2_.

**Figure 2:**
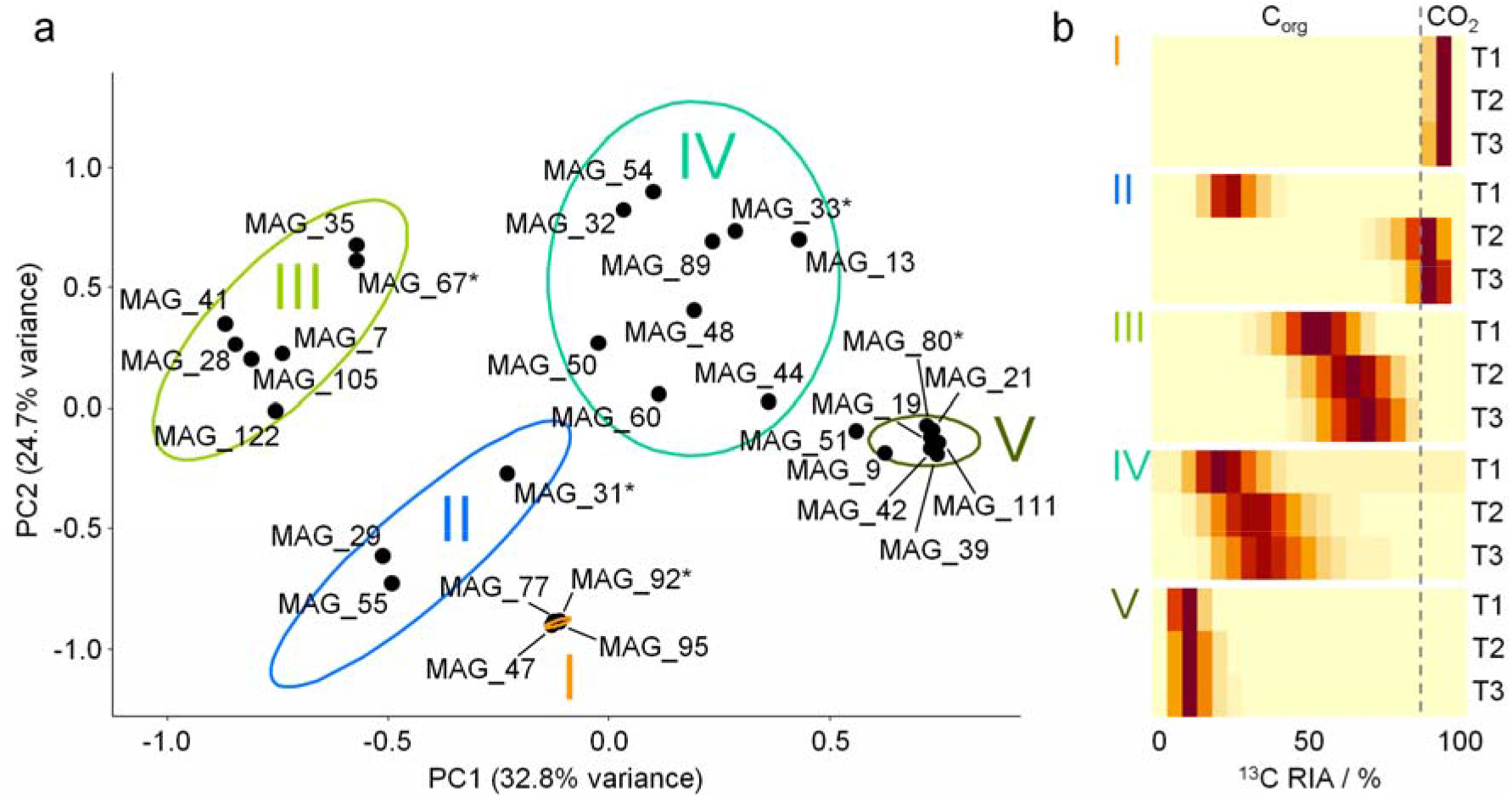
Clustering of selected MAGs based on carbon utilization. (a) Stable isotope cluster analysis based on PCA of ^13^C incorporation profiles over incubation time obtained from SIP-metaproteomics of ^13^C-microcosm samples. Each point represents a distinct organism represented by one MAG. MAG clusters are indicated by Latin numbers. Ellipses depict 95% confidence intervals. All MAGs shown facilitated the acquisition of at least two replicates of ^13^C incorporation patterns per time point. (b) Representative ^13^C incorporation profiles of MAGs marked with asterisks are given for each cluster. Heatmaps depict the extent of ^13^C incorporation in peptides of the corresponding MAG after 21 (T1), 43 (T2), and 70 days (T3) of incubation (5% intervals, ranging from 0 to 100% ^13^C relative isotope abundance).

**Figure 3:**
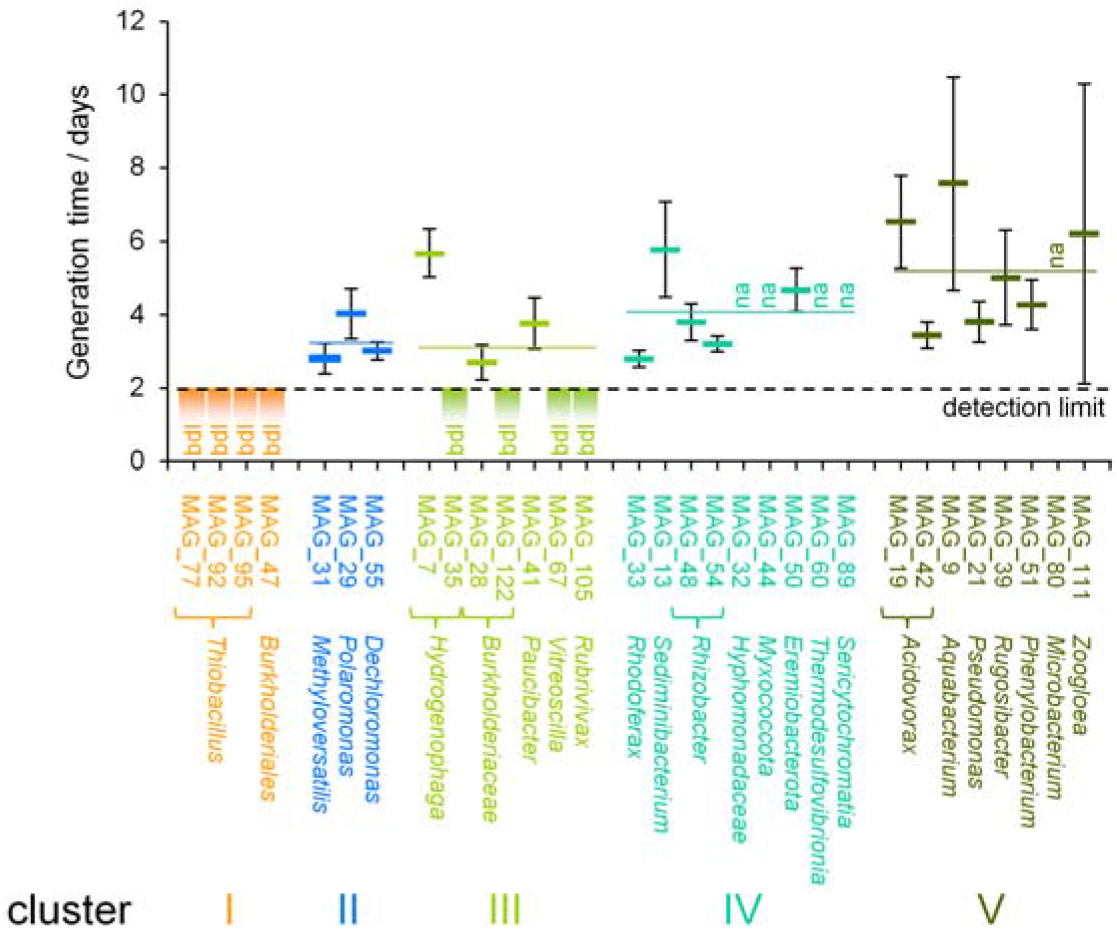
Generation times of groundwater microorganisms. Values were determined for the first 3 weeks of incubation, based on the relative abundance of ^12^C and ^13^C peptides. Shown are mean and standard deviation based on n ≥ 4 replicate determinations. Colored horizontal lines indicate average generation time for each cluster. bdl: generation time fell below the detection limit of 2 days. na: quantification of generation time was not possible.

Organisms represented by MAGs in cluster II were most closely related to species of *Methyloversatilis*, *Polaromonas* and *Dechloromonas* (all *Burkholderiales*). These microbes exhibited a moderate 65% ^13^C RIA after 21 days of incubation (Fig. 2), which suggested the utilization of labeled organic carbon from primary production as well as unlabeled organic carbon from the groundwater. After incubating for 43 and 70 days, however, ^13^C RIA increased to 91% (*p* = 1.573 × 10^−3^, *t* = −3.5225, *df* = 26.464, two-sided Welch’s *t*-test), indicative of a switch to chemolithoautotrophic growth as organic carbon became limited. Exhibiting generation times between 2 and 4 days (Fig. 3), MAGs representing these mixotrophs were more than twice as abundant as those of cluster I, accounting for 26% of the total normalized coverage.

Over the first 21 days of incubation, mean ^13^C RIAs of cluster III and IV microbes increased from 65 to 76% and from 18 to 53%, respectively (*p* = 2.211 × 10^−13^, *t* = −8.4984, *df* = 97.694 [cluster III] and *p* < 2.2 × 10^−16^, *t* = −11.626, *df* = 58.764 [cluster IV], two-sided Welch’s *t*-test; Fig. 2). This increasing trend of ^13^C RIA demonstrated two important points: First, it clearly indicated heterotrophic growth, based in part on organic carbon produced by chemolithoautotrophic organisms of clusters I and II. Second, it illustrated the increased labeling of available organic carbon, through the fixation of ^13^CO_2_. Variations observed in ^13^C RIAs between species suggested different extents of cross-feeding on chemolithoautotrophically produced organic carbon, potentially due to preferences for different organic carbon compounds. Cluster III housed the largest fraction of the MAG population, accounting for 28% of the total normalized coverage, while cluster IV accounted for 20% of this total. The vast majority of organisms represented by MAGs in these clusters exhibited generation times between 3 and 4 days (Fig. 3). However, cluster III microbes most closely related to species of *Hydrogenophaga*, *Vitreoscilla*, and *Rubrivivax* exhibited growth rates as fast as their cluster I counterparts.

In cluster V, average ^13^C RIAs reached 6% after 21 days of incubation and did not change thereafter, which hinted at active heterotrophic lifestyles early on in the experiment. Nonetheless, these organisms represented 15% of the total normalized coverage of all clusters. Generation times for cluster V microbes were slightly longer and more variable, ranging from 3.5 days for species of *Acidovorax* to eight days for *Aquabacterium* spp. (Fig. 3).

Analyses of corresponding peptide RIAs of all analyzed MAGs showed that 43, 68, and 80% of all carbon available to the microbial population was replaced with ^13^C following 21, 43, and 70 days of incubation, respectively. Quantitative DNA-SIP confirmed this labeling pattern via increases in the number of, and buoyant density shifts associated with, ^13^C-labeled OTUs (Fig. S6; Supp. Info.). SIsCA revealed carbon transfer from autotrophic cluster I to mixotrophic cluster II, and from these two further to the heterotrophs of cluster III through V through cross-feeding on ^13^CO-derived organic carbon.

### Functional characterization of MAGs reveals putative mixotrophs

All of the putative autotrophs detected employed the Calvin-Benson-Bassham (CBB) cycle for CO_2_ fixation (Fig. 4). Subunits of the enzyme ribulose-1,5-bisphosphate carboxylase/oxygenase (RuBisCO) were detected in the proteomes of 15 of 31 MAGs, and 14 of these contained additional enzymes of the CBB cycle. No other complete CO_2_ fixation pathways were identified. Proteins of the CBB cycle were present not only in strict or facultative autotrophs of cluster I (i.e., relatives of *Thiobacillus* spp.) or cluster II (e.g., relatives of *Methyloversatilis, Polaromonas*, and *Dechloromonas* spp.), but also in heterotrophic organisms most closely related to species of *Hydrogenophaga*, *Rhodoferax*, *Paucibacter*, and *Rubrivivax* of clusters III and IV. Mixotrophs comprised > 50% of all microbial taxa represented across all clusters, which underscored the immense importance of their contributions to carbon cycling in modern groundwater. The mixotrophic lifestyle, by no means a rare or insignificant trait in these groundwater microcosms, appeared to bestow fitness on the microbes.

**Figure 4:**
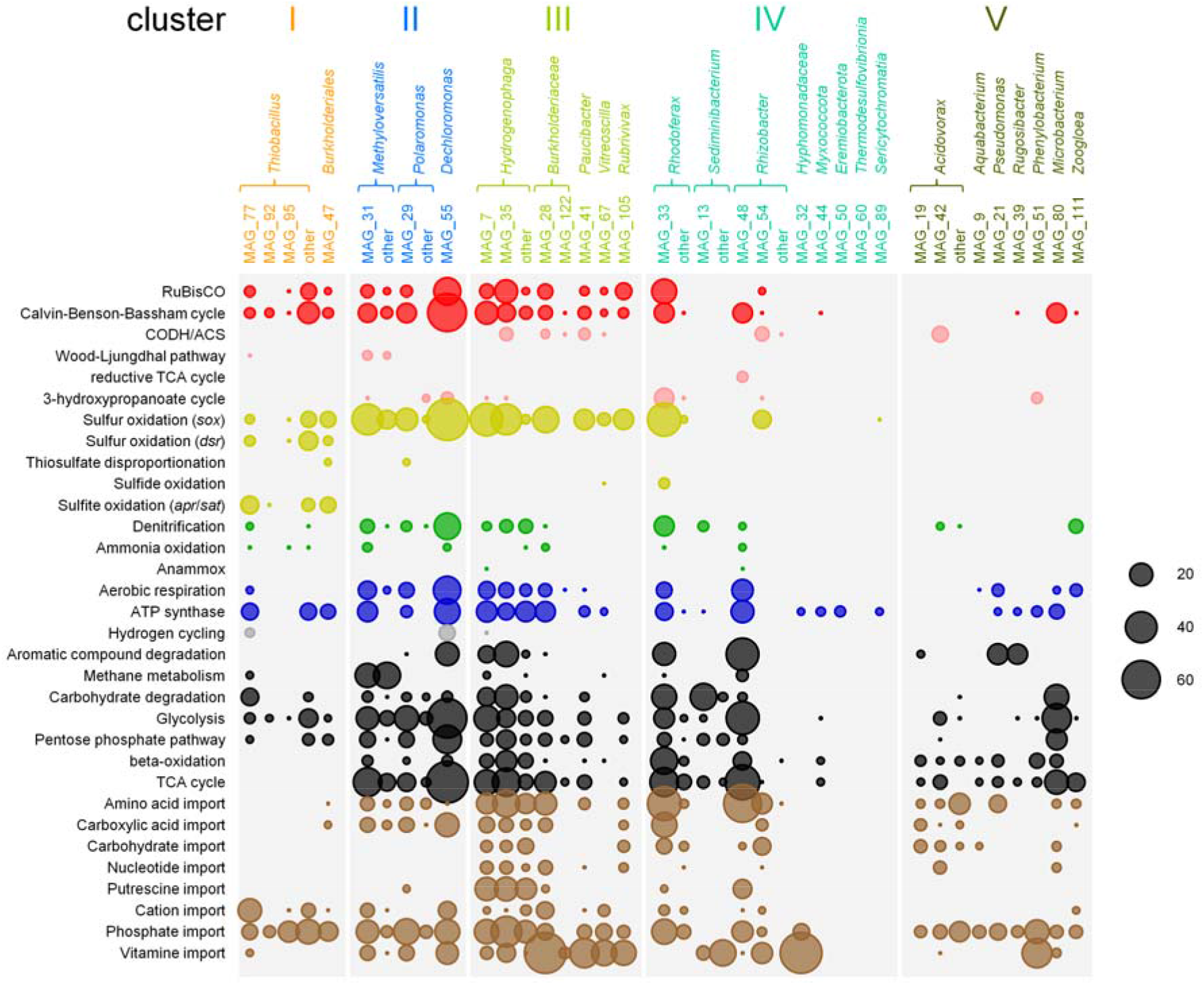
Metabolic functionality of selected MAGs. The sizes of the bubbles correspond to the total number of peptides detected for each MAG and each functional category identified at any time point. Metabolic functions are grouped into CO_2_ fixation (red), sulfur cycling (yellow), nitrogen cycling (green), aerobic respiration and ATP synthesis (blue), organic carbon utilization (black), and import functions (brown). The taxonomic categories “other” include peptides that were assigned to multiple MAGs affiliated with the same genus. Only MAGs considered in the stable isotope cluster analysis are shown. RuBisCO: ribulose-1,5-bisphosphate carboxylase/oxygenase, CODH/ACS: carbon monoxide dehydrogenase/acetyl-CoA synthase, TCA cycle: tricarboxylic acid cycle.

### MAGs express pathways for the utilization of reduced sulfur compounds

Sixteen MAGs expressed proteins for sulfur oxidation via the Sox or Dsr enzyme system (Fig. 4). Cluster II, III, and IV microbes phylogenetically affiliated with species of *Methyloversatilis*, *Dechloromonas*, *Hydrogenophaga*, *Rhodoferax* and other *Betaproteobacteriales* utilized the Sox system exclusively. MAGs harbored gene clusters of the conserved soxCDYZAXB gene order (Fig. S7), featuring the core components of the Kelly-Friedrich pathway^53,54^. This pathway facilitates the complete oxidation of thiosulfate to sulfate, without free intermediates^29^. Accessory genes *soxVW*, *soxEF*, *soxTRS*, and *soxH* were randomly distributed through the MAGs disconnected from the main operon.

Cluster I microbes most closely related to *Thiobacillus* spp. produced enzymes for both the Sox and Dsr system, and corresponding MAGs housed a truncated *soxXYZAB* gene cluster that lacked genes *soxCD* required to oxidize the sulfane group of thiosulfate. As such, these organisms likely used the branched thiosulfate oxidation pathway typical for *Thiobacillus* spp.^55^, whereby Dsr operating in reverse oxidizes the sulfane-derived sulfur atom to sulfite, with elemental sulfur as intermediate^29^. Cluster I MAGs maintained the conserved operon structure *dsrABEFHCMKLJOPNR*, including genes *dsrEFH* and *dsrL* typical for sulfur oxidizers but lacking gene *dsrD* for sulfate reduction^28^. These organisms also expressed *aprAB* and *sat*, which encode Adenosine-5’-phosphosulfate reductase and ATP sulfurylase, respectively, each of which can function in reverse to oxidize sulfite to sulfate ^56^. Hence, groundwater facultative chemolithoautotrophs employed the Sox system to oxidize thiosulfate to sulfate, while obligate chemolithoautotrophs utilized an incomplete version of this system to oxidize the sulfone group and the Dsr/Apr/Sat system to oxidize the sulfane group of thiosulfate.

### Use of alternative electron acceptors and donors in sulfur oxidizers

Cytochrome c oxidase and other enzymes of the respiratory chain were detected in 15 sulfur oxidizer MAGs, 12 of which also harbored enzymes for nitrate reduction (i.e., nitrate reductase, nitrite reductase, nitric oxide reductase; Fig. 4). Several sulfur oxidizers related to species of *Dechloromonas* and *Rhodoferax* expressed both pathways concurrently. Proteins for ammonia oxidation (i.e., ammonia monooxygenase, hydroxylamine oxidoreductase) were produced by a variety of cluster I and IV microbes, such as *Thiobacillus* and *Methyloversatilis* species. MAG_77 (*Thiobacillus*), MAG_55 (Dechloromonas) and MAG_7 (Hydrogenophaga) even expressed [NiFe]-hydrogenase genes.

### Utilization of organic carbon in oligotrophic groundwater

Cluster I, II, and III MAGs exhibited a gradient of increased versatility in utilizing various organic carbon compounds. While cluster I’s strict autotrophs only expressed pathways for sugar degradation, MAGS of clusters II through V produced proteins germane to the breakdown and transport of simple sugars (e.g., glycolysis, pentose phosphate pathway), amino acids (TCA cycle), fatty acids (beta-oxidation), C_1_ compounds, and aromatics (Fig. 4). The TCA cycle was one of the most abundant metabolic modules observed in MAGs of cluster II to V. Degradation pathways for toluene and ethylbenzene were expressed by organisms most closely related to species of *Dechloromonas* and *Rhizobacter* (*Betaproteobacteriales*), respectively. Enzymes for naphthalene and catechol catabolism were detected in MAGs representing organisms related to *Hydrogenophaga* and Pseudomonas spp., while gene products germane to the degradation of complex carbohydrates (e.g., starch, chitin) were produced by MAGs representing relatives of *Microbacterium* and *Sediminibacterium* species. The metabolic machinery required to metabolize C_1_ compounds was detected primarily in microbes related to *Methyloversatilis* spp., which typically possessed methanol dehydrogenase, formate dehydrogenase, and other enzymes involved in tetrahydromethanopterin-dependent C_1_-cycling.

Gene products relevant to import systems for amino acids and carboxylic acids (e.g., alpha-keto acids, C_4_-dicarboxylates, lactate) were overly abundant in mixotrophs and heterotrophs of clusters II to V (Fig. 4). Cluster III to V microorganisms that had grown exclusively heterotrophically exhibited the greatest diversity of import-related proteins, including those for the transport of carbohydrates and nucleotides. Only transporters targeting cations (predominantly iron) and phosphate were detected in MAGS representing cluster I obligate autotrophs.

## Discussion

Despite conditions strongly favoring autotrophic sulfur oxidizers, mixotrophs – not obligate chemolithoautotrophs, were the most abundant active microorganisms in the groundwater microcosms. While a diverse microbial consortium was detected, strict chemolithoautotrophs accounted for only 3% of the total groundwater biodiversity. This is astonishing since thiosulfate and oxygen were readily available throughout the experiment, which should have selectively promoted the proliferation of chemolithoautotrophic microbes. Genome-resolved SIP-Metaproteomics combined with our novel SIsCA approach facilitated identification of active microbes, characterization of their expressed gene products (and linked pathways), and quantification of carbon uptake and transfer within a diverse community over time (Fig. 5). Furthermore, highly sensitive Raman spectroscopy showed that microbes were active at the outset of the incubation (no discernable lag phase), despite low sulfur oxidation rates. Shedding new light on the mechanisms by which CO_2_-derived carbon is assimilated and cycled by groundwater microflora, this approach far surpasses others that have implicated the importance of chemolithoautotrophy in groundwater based solely on functional gene and metagenomics data^13,14,17,57^. Within 21 and 70 days of incubation, 43 and 80% of the total groundwater biomass consisted of CO_2_-derived carbon, respectively. It is convincingly clear that this rapid enrichment of CO_2_-derived carbon did not occur in fixed, linear progression from chemolithoautotrophs to heterotrophs, but through a highly complex and reticulated web of trophic interactions dominated by mixotrophs – the experts of organic carbon recycling.

**Figure 5:**
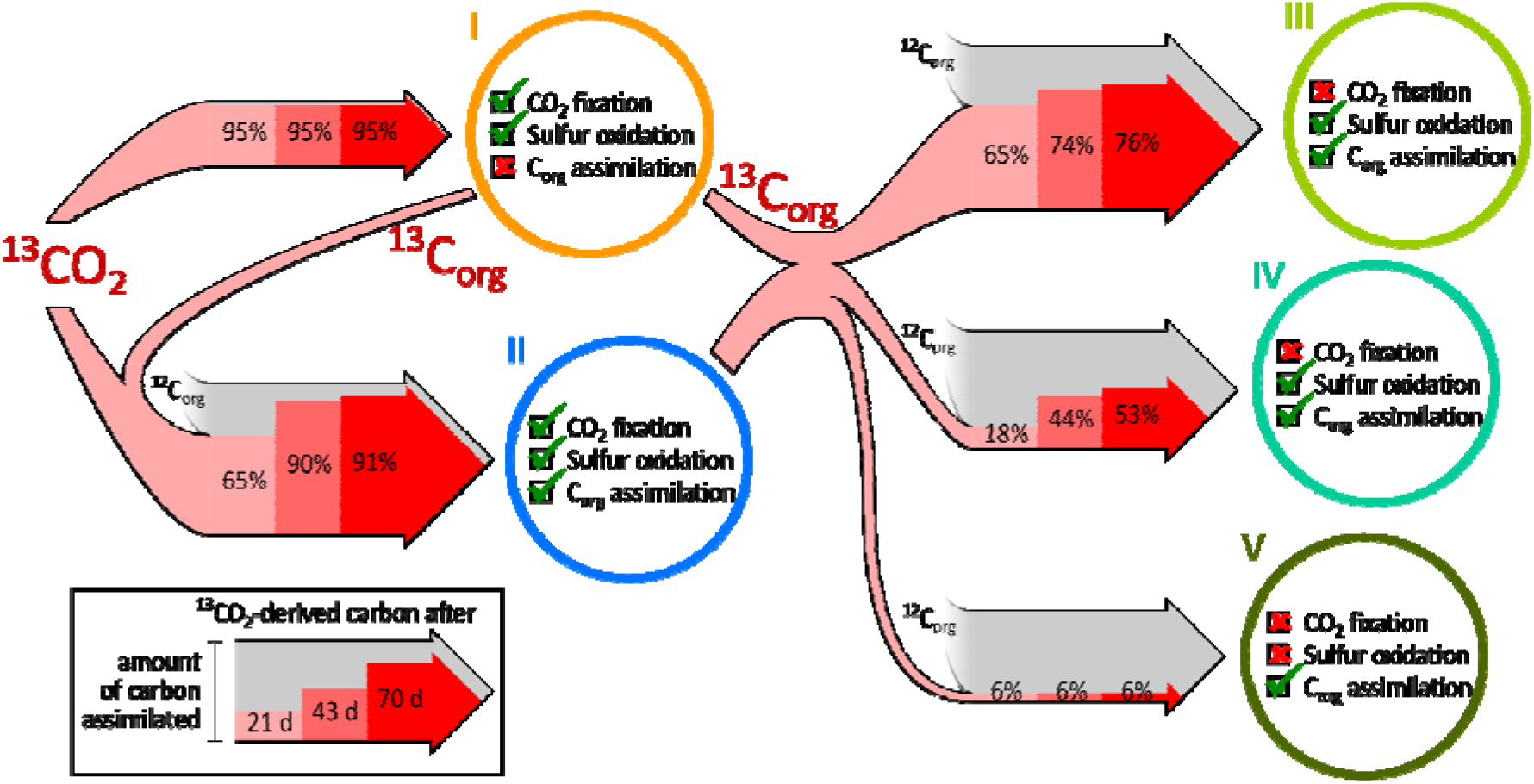
Carbon flux between microbial clusters. Red arrow inlays illustrate the fraction of ^13^CO - derived carbon assimilated by each microbial cluster after 21, 43, and 70 days. Arrow width scales with the total amount of carbon assimilated based on the relative abundance of the respective microbial cluster in the metagenomics analysis. Fading grey arrows indicate uptake of unlabeled organic carbon from the groundwater. Checkboxes highlight the presence and activity of metabolic functions for CO_2_ fixation, utilization of organic carbon, and sulfur oxidation.

These mixotrophs strongly preferred heterotrophic growth to the fixation of CO_2_, presumably a consequence of the greater metabolic cost of carbon assimilation via the CBB cycle^58,59^. The ability to fix CO_2_ affords these microbes the luxury of an opportunistically selective lifestyle, which lends itself to bolstered fitness (and rapid dominance) when organic carbon becomes limited in oligotrophic systems. Cluster II mixotrophs, for example, transitioned from heterotrophy to CO_2_ fixation late in the incubation, likely due to such limitations. In a similar vein, cluster III mixotrophs expressed pathways for autotrophic growth but were never required to fix CO_2_. These microbes were able to access a more diverse repertoire of carbon sources due to a greater metabolic versatility in organic carbon utilization.

In support of the higher fitness associated with the opportunistic CO_2_ fixation, mixotrophs grew considerably faster (generation times of two days or less) than cluster IV and V organisms restricted to an exclusively heterotrophic lifestyle (generation times up to 8 days). Surprisingly, however, these heterotrophs were also able to oxidize reduced sulfur compounds, suggestive of a chemolithoheterotrophic lifestyle. With respect to energy conservation, the constant influx of reduced sulfur via weathering of interspersed pyrite minerals^25–27^ renders sulfur oxidation an attractive alternative to the oxidation of organic compounds, both in Hainich CZE groundwaters and beyond. As the energetic requirement for CO_2_ fixation is greater than the potential gain from organic carbon oxidation, the most efficient strategy for both mixotrophs and heterotrophs is to net the greatest amount of possible from sulfur oxidation and preserve precious organic carbon for anabolic demands.

The diversity of organic carbon utilization motifs shifted gradually, and inversely, with CO_2_ fixation. At one end of the transition were the strict autotrophs of cluster I, relying exclusively on CO_2_ as carbon source. No organic carbon transporters were detected for any of these organisms. Their limited metabolic breadth restricted growth to that from simple sugars, likely to utilize carbon assimilated via the CBB cycle^58^. At the other end of the transition were organisms from clusters IV and V that assimilated organic carbon exclusively. To endure the groundwater environment *sans* autotrophic CO_2_ fixation machinery, these organisms had to maintain and express a wide variety of organic carbon transport and assimilation pathways. The most fit organisms in this modern groundwater ecosystem, however, were the mixotrophs of clusters II and III. Establishing dominance by opportunistically exploiting their physiological flexibility, these organisms rapidly outcompeted their strictly autotrophic brethren (5-fold greater abundance).

Organisms most closely related to Burkholderiales spp., the key mixotrophic taxa in our groundwater microcosms, gave rise to the greatest number of RuBisCO-encoding transcripts in a previous study at our groundwater site^17^. For taxa like *Polaromonas*, *Dechloromonas*, *Hydrogenophaga*, and Rhodoferax spp., the ability to oxidize sulfur has been posited based solely on genomic evidence^60–63^. Hitherto, chemolithoautotrophic growth on reduced sulfur compounds has not been observed from any of these genera in pure culture. Our study demonstrates that these organisms can use reduced sulfur as an energy source, and species of *Polaromonas*, *Dechloromonas*, and potentially **Hydrogenophaga** used it to fuel autotrophic growth. These sulfur oxidizers expressed pathways for both aerobic respiration and denitrification, despite the fact that no nitrate was added and nitrate concentrations in the groundwater of this well never exceeded 10 mg/L^27^. Constitutive maintenance and expression of denitrification enzymes is likely more energetically cost effective than regulating gene expression^64^. This strategy also affords these microbes the advantage of utilizing different electron acceptors when oxygen becomes limited.

Utilizing an incomplete TCA cycle that precludes heterotrophic growth, the *Thiobacillus*-related organisms of cluster I are known to be obligately autotrophic^65^. Previously, by carrying out thiosulfate- and hydrogen-driven denitrification, *Thiobacillus* spp. grew up to represent upwards of 50% of an enrichment culture obtained from Hainich CZE groundwater^66^. In situ, however, *Thiobacillus* spp. are typically found in lower numbers^17^, and most commonly appear in deeper, more CO-rich subsurface systems^13^. This suggests diminished fitness and inability to compete with more physiologically fit mixotrophs in oligotrophic modern groundwater. *Thiobacillus* can store the elemental sulfur produced as intermediate by the Dsr enzyme system in periplasmic granules^65,67^. This storage might allow the organism to withstand times where no reduced sulfur compounds in the groundwater are available.

There are two key advantages to being a mixotrophic sulfur oxidizer in the groundwater habitat. First and foremost, these cells exist completely independent of surface carbon input dynamics. The energy sources they rely on is produced autochthonously in the geological setting. Second, their diverse breadth of physiological capabilities allows these microbes to modulate the means by which they satisfy their anabolic requirements and energy demands based on the types of carbon available. This includes carbohydrate degradation pathways for surface-derived plant polymers^8,68^, amino acid and nucleotide uptake systems for microbially-derived carbon^69,70^, C_1_ metabolic functions for C_1_ carbon compounds from biomass degradation^71^, and hydrocarbon degradation pathways for rock-derived carbon^72,73^. We hypothesize that similar strategies exploiting a myriad of carbon assimilation pathways and versatile energy acquisition motifs benefit microbes dominating other oligotrophic systems, such as boreal lakes or the upper ocean^74,75^.

## Conclusions

Our novel SIsCA-based approach facilitated the quantitative and temporal resolution of carbon flux through key subpopulations of a modern groundwater microbiome. Mixotrophs dominated this oligotrophic environment by fulfilling and supplementing their organic carbon requirements via opportunistic fixation of CO_2_. This CO_2_-derived organic carbon was rapidly incorporated into, and recycled throughout, microbial biomass through a highly efficient and complex trophic network. To mitigate low levels of organic carbon, autotrophic, mixotrophic, and heterotrophic microorganisms utilized reduced sulfur compounds as energy sources and preserved what organic carbon was available for anabolic demands. A wide variety of carbon assimilation pathways enabled mixotrophs and heterotrophs to make optimal use of the scarce amounts of organic carbon characteristic of oligotrophic environments. We posit that the concerted, opportunistic deployment of a wide variety of highly versatile pathways for assimilating carbon and generating energy from inorganic sources is key to microbial success in oligotrophic environments. The findings of this investigation significantly enhance our understanding of microbial survival strategies and their role in ecosystem functioning while demonstrating the powerful utility of next-generation physiology approaches like SIsCA in testing hypotheses established in metagenomics-based endeavors.

## Declarations

### Ethics approval and consent to participate

Not applicable

### Consent for publication

Not applicable

### Availability of data and materials

Metagenomic and amplicon sequencing data that support the findings of this study have been deposited into NCBI under the BioProject accession PRJNA633367. Mass spectrometry proteomics data have been deposited into the ProteomeXchange Consortium via the PRIDE^76^ partner repository with the dataset identifier PXD024889.

### Authors’ contributions

MT and KK conceived and designed the study. MT conducted the microcosm experiments and molecular biology work. GAM, RH, PR, and JP conducted the Raman microspectroscopic analyses. NJ and MvB conducted mass spectrometric analysis for metaproteomics. MT analyzed the metagenomics data with the assistance of WAO and BMH, and analyzed the SIP-metaproteomics data. MT wrote the manuscript with contributions from all authors.

## Acknowledgments

We are grateful to Robert Lehmann, Falko Gutmann, Heiko Minkmar, Jens Wurlitzer, and Lena Carstens for assistance with field and lab work, and sampling of groundwater. We thank Julian Hniopek for help with Raman measurements, and Daniel Desirò and Martin Hölzer for assistance with sequencing. We also extend gratitude to Ivonne Görlich and Marco Groth at the Core Facility DNA sequencing of the Leibniz Institute on Aging - Fritz Lipmann Institute in Jena for help with Illumina sequencing.

