## Supplementary Information for "Bolstering fitness via opportunistic CO_2_ fixation: mixotroph dominance in modern groundwater"

Short title: Chemolithoautotrophy in shallow groundwater

### Supplementary Methods

#### Amplicon sequencing

For taxonomic characterization of the bacterial community in the microcosms, amplicon sequencing of the bacterial 16S rRNA gene, region V3 to V5, was done. Polymerase chain reaction was performed using primer pair Bact_341F/Bact_805R^1^ and HotStarTaq Mastermix (Qiagen, Hilden, Germany) as described previously^2^. Amplicons were purified using NucleoSpin Gel & PCR Clean-Up Kit (Macherey-Nagel, Düren, Germany). The NEBNext Ultra DNA Library Prep Kit for Illumina (New England Biolabs, Frankfurt, Germany) was used to prepare libraries for amplicon sequencing, following the manufacturer’s instructions. Amplicons were purified using AMPure XP Beads (Beckman Coulter, Krefeld, Germany), and amplicon sequencing was then carried out in-house on a MiSeq Illumina platform (Illumina, Eindhoven, The Netherlands) with v3 chemistry.

Raw sequence data was analyzed using mothur (v.1.39)^3^, according to the mothur standard operating procedures^4^ as previously described^5^. OTU binning with a 3% identity cutoff was performed, followed by OTU classification using the SILVA reference database release SSU 132^6^. Raw Illumina MiSeq sequencing data have been deposited in the Sequence Read Archive (SRA) of NCBI under BioProject accession PRJNA633367.

#### Assessment of DNA density shifts

To evaluate the ^13^C incorporation on nucleic acid level, DNA samples extracted from the triplicate groundwater microcosms supplemented with ^13^C and ^12^C bicarbonate after 21 and 70 days of incubation were subjected to SIP ultracentrifugation in CsCl gradients as previously described^7^. Ultracentrifugation was carried out in an NVT 90 rotor (Beckman Coulter, Brea, CA, USA) in a Sorvall Discovery 90SE ultracentrifuge (Thermo Fisher Scientific, Waltham, MA, USA) at 40,900 rpm and 20 °C for 60 h. The gradients were separated into 12 to 14 fractions, covering a buoyant density range from 1.77 g ml^-1^ to 1.68 g ml^-1^. The buoyant density of each fraction was determined using a Reichert AR200 digital refractometer (Reichert Analytical Instruments, Depew, NY, USA). To account for tube-to-tube and spin-to-spin variations of the density gradient^8^, gradients were normalized by centering the density of ^12^C DNA onto 1.70 g ml^-1^. The DNA in the density fractions was purified by precipitation with NaCl-PEG as previously described^9^ and quantified fluorometrically using the Qubit dsDNA broad-range assays (Thermo Fisher Scientific). Subsequently, amplicon sequencing of the bacterial 16S rRNA gene was performed as described above. OTU-wise DNA buoyant density profiles over the gradients were obtained as previously described^10^. Only OTUs that were represented by at least 10 reads in one fraction of each ^12^C and each ^13^C replicate were included in the analysis. This was done separately for both time points. From the DNA buoyant density profiles of each OTU, the DNA density of that OTU ($\overline{\rho}_{OTU}$) was determined by performing least-squares regression of the OTU abundance $A_{OTU}$ per fraction $f$ to the fraction density $\rho_{f}$ using a nonlinear model (I), where $\alpha^{2}$ represents the variance of the OTU DNA density.

$A_{OTU,f}=e^{-\frac{\left( \rho_{f}-\overline{\rho}_{OTU} \right)^{2}}{2\alpha^{2}}}$	(I)

The average ^12^C and ^13^C OTU DNA density was determined as the mean of the respective triplicates. The density shift between ^12^C and ^13^C samples was determined separately for each time point. The significance of the shift was assessed using Student’s *t*-test based on the respective triplicates.

### Supplementary Results

#### Functional composition of the whole microbial community

Mapping the functional information obtained by SIsCA and genome-resolved metaproteomics to the corresponding taxa in 16S rRNA gene profiles allowed us to classify the lifestyle of up to 50% of the microbial community (Figure S4). Over all time points, only 3.2 ± 3.1% (mean ± sd) of the community were composed of strict autotrophs, primarily affiliated with *Thiobacillus*. Mixotrophs comprised 17.6 ± 4.3% of the total community and were dominated by *Rhodoferax* and *Hydrogenophaga*, but the largest fraction of the total community, with 20.1 ± 7.8%, consisted of heterotrophs, primarily affiliated to *Sediminibacterium* (*Bacteroidetes*), *Pseudomonas* (*Gammaproteobacteria)*, *Sericytochromatia* (*Cyanobacteria*) and *Microbacterium* (*Actinobacteria*) (Figure S5).

#### DNA-SIP supports role of ^13^CO_2_-derived carbon

In addition to the Stable Isotope Cluster Analysis (SIsCA) approach based on metaproteomics data, quantitative DNA-SIP was performed to provide a higher coverage of the microbial community. The number of ^13^C-labeled taxa increased from 21 OTUs after 21 days of incubation to 65 OTUs after 70 days of incubation, observable by a significant shift of DNA buoyant density (Figure S6). While after 21 days, these OTUs were mainly affiliated with *Burkholderiales* such as *Thiobacillus*, *Hydrogenophaga* and *Polaromonas*, after 70 days, various other *Alpha*- and *Gammaproteobacteria* were included. The average buoyant density shift of these OTUs likewise increased significantly from 0.021 ± 0.010 g ml^-1­^ to 0.028 ± 0.013 g ml^-1^ (*p* = 0.014, *t* = -2.2842, *df* = 38.264, one-sided Welch’s *t*-test) in this period. This highlights the increasing role of ^13^CO_2_-derived carbon introduced by chemolithoautotrophic activity into the microbial carbon pool in the groundwater incubations, and the flux of ^13^C through the microbial food web.

### Supplementary Figure and Table legends

**Figure S1: Hydrochemical conditions in groundwater microcosms.** (A) Mean values for oxygen, thiosulfate and sulfate concentrations determined in all ^12^C and ^13^C microcosms over incubation time are given. Linear regression curves are shown for 0 to 21 days (dashed line), 21 to 43 days (dotted line) and 43 to 70 days (solid line). (B) Rates of oxygen and thiosulfate consumption as well as sulfate production based on linear regression are summarized over three time intervals of incubation. The number of replicates is n=18 for 0 to 21 days, n=12 to 43 days and n=6 to 70 days. Error bars indicate standard deviation.

**Figure S2: Raman microspectroscopic analysis of the groundwater microbial community.** (A) Mean Raman spectra of groundwater bacteria incubated with heavy water (D_2_O) for 12 to 47 days. (B) Mean Raman spectra of groundwater bacteria incubated with H_2_O for 12 to 47 days. (C) Confusion matrix showing the results of the PCA-LDA model built for the classification of metabolically active bacterial cells in groundwater. The differentiation between deuterium labeled (D_2_O) and non-labeled (H_2_O) bacterial cells achieved an overall accuracy of 92.2%, a mean sensitivity of 92.6% and a mean specificity of 92.6%.

**Figure S3: Stable Isotope Cluster Analysis (SIsCA) of peptides assigned to MAGs.** The analysis is based on PCA of ^13^C incorporation profiles over incubation time obtained by SIP-metaproteomics of samples from the ^13^C-microcosms. Each point represents one peptide of an organism associated with a particular MAG. The color code is used to highlight peptides that belong to the same MAG.

**Figure S4: Functional composition of the groundwater microbial community.** Shown are relative abundances of autotrophs (cyan), mixotrophs (blue), and heterotrophs (orange) of the groundwater microcosms on DNA level. The relative quantification of microbial taxa is based on bacterial 16S rRNA gene amplicon sequencing data. Functional classification of microbial taxa into the functional groups is based on the results of the SIP-metaproteomics analysis.

**Figure S5: Taxonomic composition of the microbial community in the groundwater microcosms.** Shown are the 20 most abundant bacterial genera based on read abundance from MiSeq amplicon sequencing of bacterial 16S rRNA genes. Further genera are summarized in the category ‘others’. Each bar corresponds to one replicate microcosm. One replicate from the microcosms incubated for 21 days is not shown as the sequencing of the respective sample failed.

**Figure S6: Change of DNA buoyant density for bacterial OTUs in the groundwater microcosms.** Density shifts are indicative of ^13^C incorporation in DNA of the respective organism represented by the OTU after 21 days (grey) or 70 days (red) of incubation. Shifts were calculated as difference between density in triplicate ^12^C samples and density in triplicate ^13^C samples. Error bars represent standard deviation of the density shift. Only OTUs with significant density shifts are shown. OTUs are sorted by decreasing density shift individually for each time point. The solid lines indicate the average density shift of the shown OTUs, the dashed lines indicate the respective standard deviation.

**Figure S7: Gene clusters involved in sulfur oxidation observed in MAGs obtained from the groundwater microcosms.** (A) Clusters of *sox* genes observed in MAGs related to *Thiobacillus* showing a canonical *soxXYZAB* gene order. The *sox* cluster of *Thiobacillus denitrificans* ATCC 25259 (NC_007404.1) is given as reference. (B) Clusters of *sox* genes observed in other MAGs showing a canonical *soxCDYZAXB* gene order. The *sox* cluster of *Dechloromonas aromatica* RCB (CP000089.1) is given as reference. (C) Clusters of *dsr* genes observed in the MAGs. The *dsr* gene cluster of *Thiobacillus denitrificans* ATCC 25259 is given as reference. Genes are represented by arrows. Grey arrows show unspecific genes. For arrows with a red outline, products of the corresponding genes have been detected by metaproteomics analysis. Numbers above the arrows indicate the respective contigs of the MAGs or the gene accession numbers for references. Lines between arrows indicate gaps larger than 100 nucleotides. Ellipses (‘…’) indicate splits between contigs or gaps of more than 5,000 nucleotides on one contig. Scale bar depicts gene length of 1,000 nucleotides. Time point and replicate of the sample the respective MAG was obtained from is shown below the MAG description.

**Table S1: Accession numbers and taxonomic affiliation of metagenome-assembled genomes (MAGs).**

**Figure S1:**

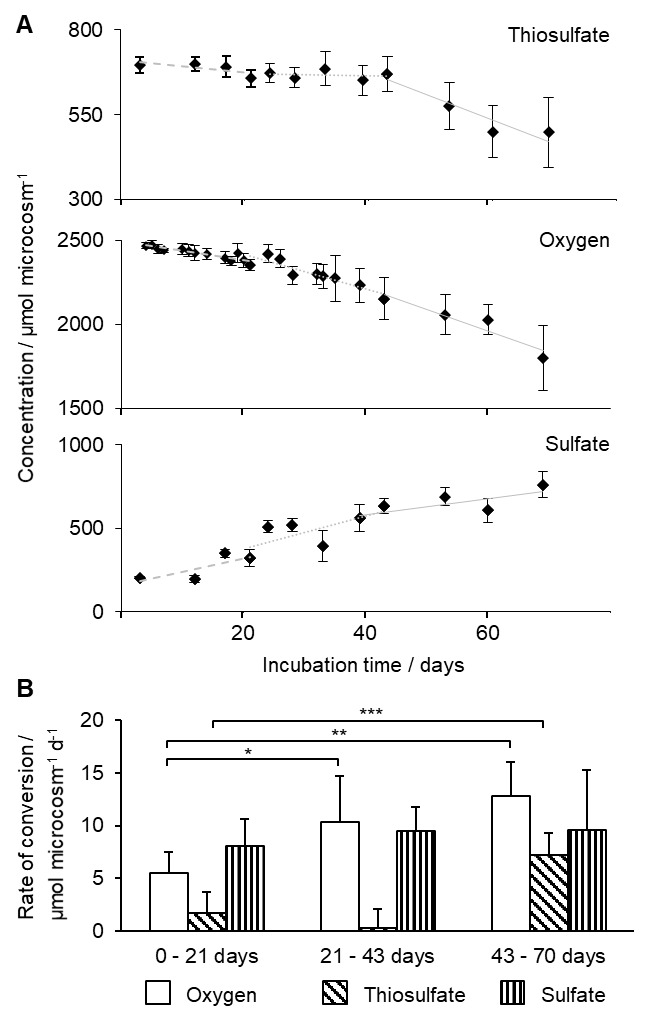

**Figure S2:**

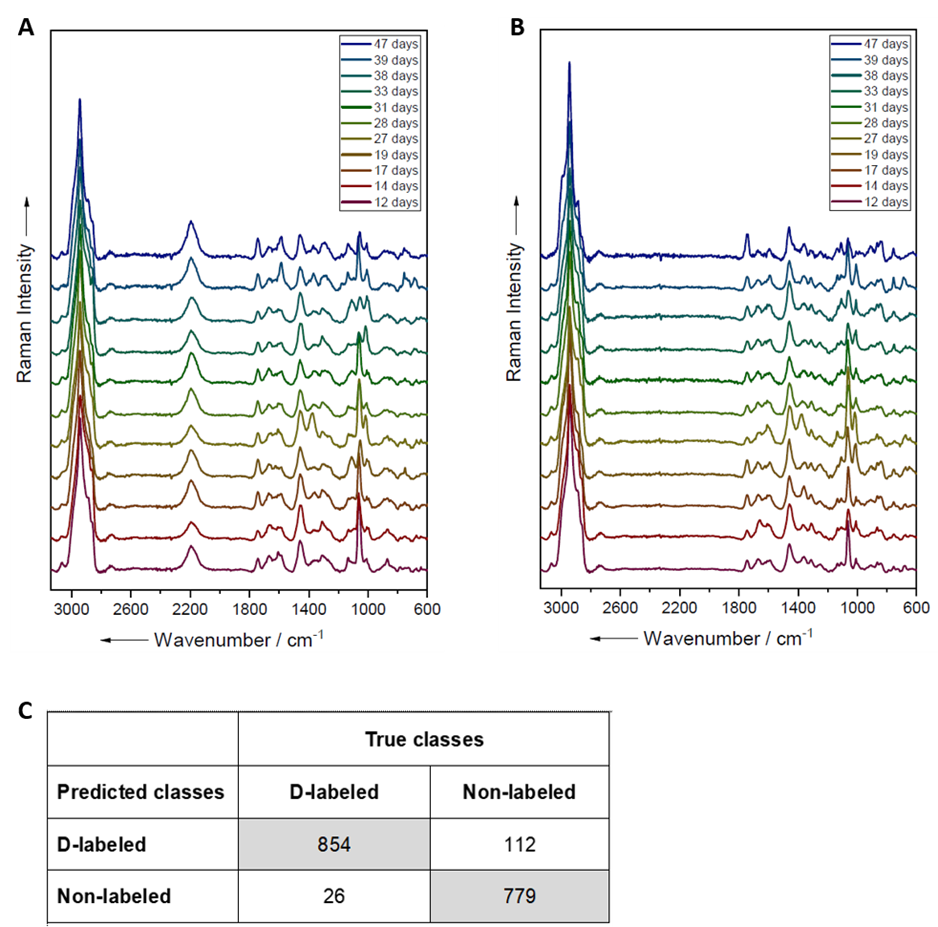

**Figure S3:**

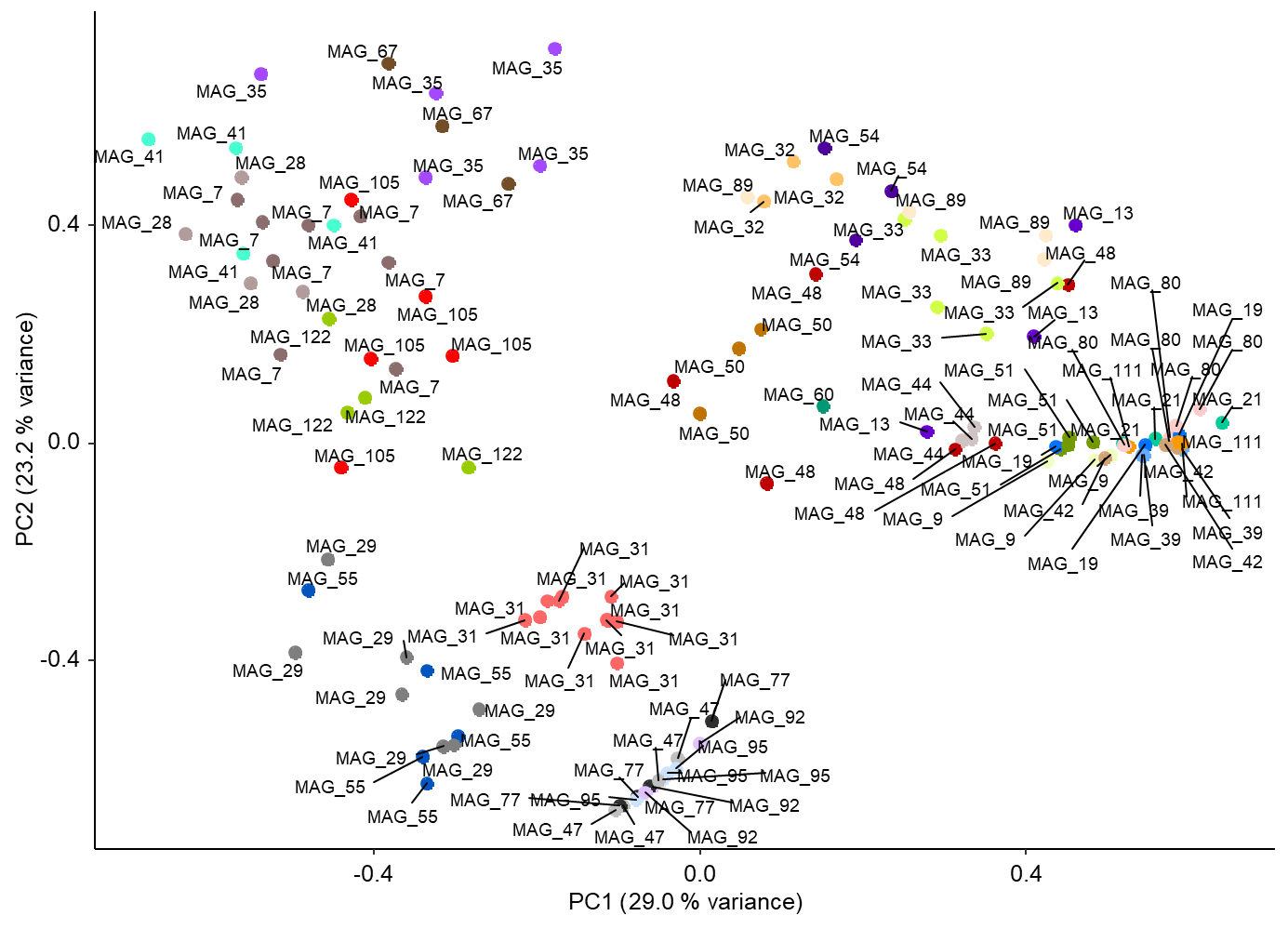

**Figure S4:**

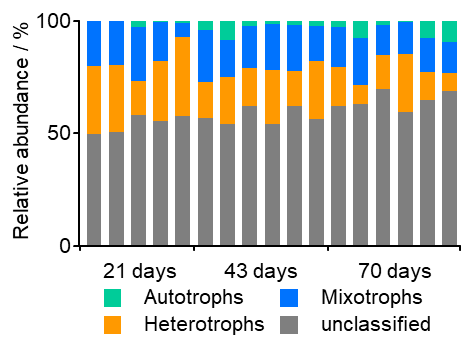

**Figure S5:**

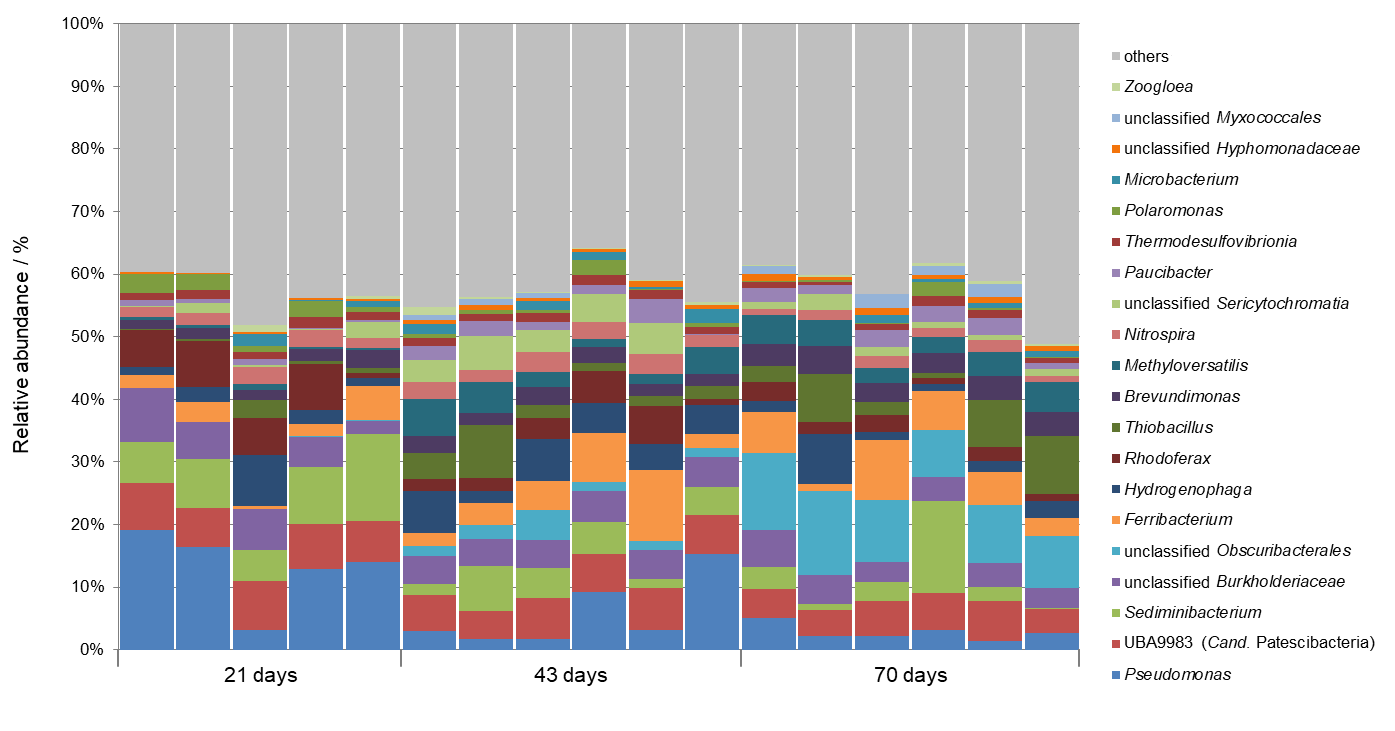

**Figure S6:**

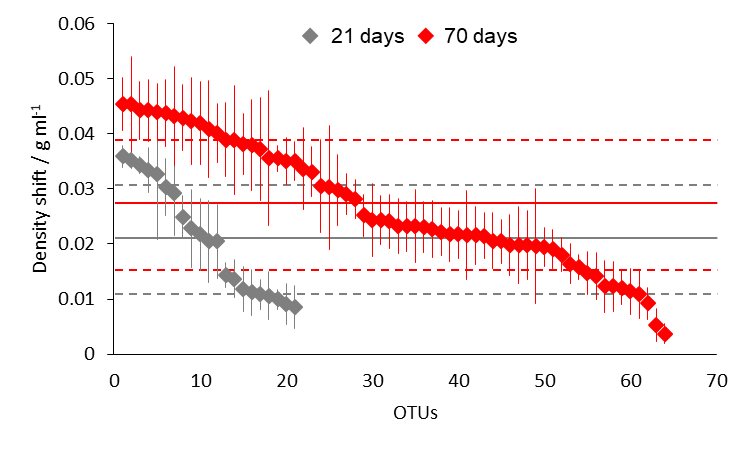

**Figure S7:**

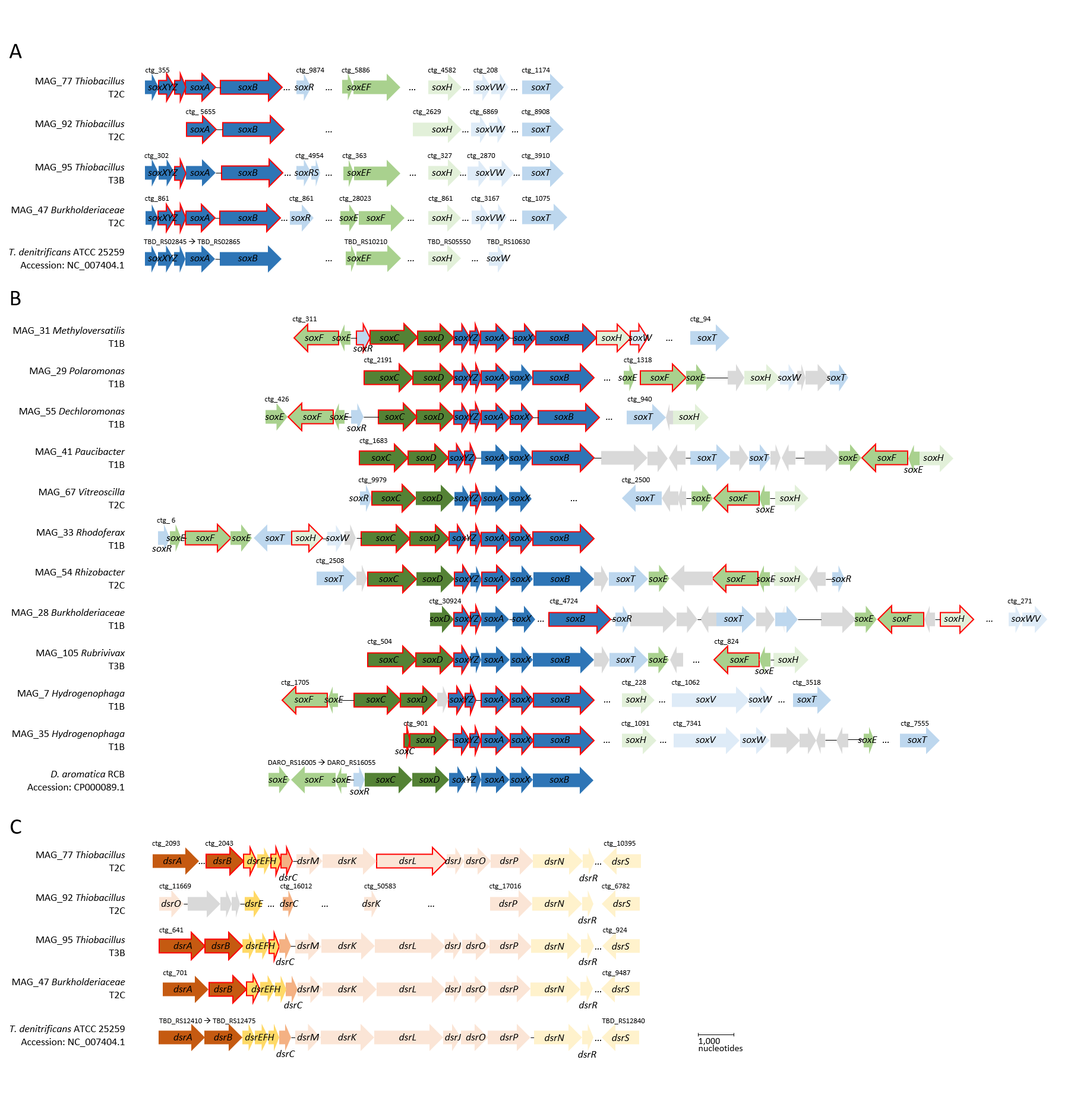

**Table S1:**

| BioSample | Genome Accession | Genome name | Taxonomy |
| --- | --- | --- | --- |
| SAMN16635724 | JADMJG000000000 | MAG_13 | *Sediminibacterium* sp. |
| SAMN16635725 | JADMJH000000000 | MAG_19 | *Acidovorax* sp. |
| SAMN16635726 | JADMJI000000000 | MAG_21 | *Pseudomonas* sp. |
| SAMN16635727 | JADMJJ000000000 | MAG_28 | Burkholderiaceae bacterium |
| SAMN16635728 | JADMJK000000000 | MAG_29 | *Polaromonas* sp. |
| SAMN16635729 | JADMJL000000000 | MAG_31 | *Methyloversatilis discipulorum* |
| SAMN16635730 | JADMJM000000000 | MAG_32 | Hyphomonadaceae bacterium |
| SAMN16635731 | JADMJN000000000 | MAG_33 | *Rhodoferax* sp. |
| SAMN16635732 | JADMJO000000000 | MAG_35 | *Hydrogenophaga* sp. |
| SAMN16635733 | JADMJP000000000 | MAG_39 | *Rugosibacter* sp. |
| SAMN16635734 | JADMJQ000000000 | MAG_41 | *Paucibacter* sp. |
| SAMN16635735 | JADMJR000000000 | MAG_42 | *Acidovorax* sp. |
| SAMN16635736 | JADMJS000000000 | MAG_44 | Myxococcales bacterium |
| SAMN16635737 | JADMJT000000000 | MAG_47 | Burkholderiales bacterium |
| SAMN16635738 | JADMJU000000000 | MAG_48 | *Pseudomonas umsongensis* |
| SAMN16635739 | JADMJV000000000 | MAG_50 | Eremiobacterota bacterium |
| SAMN16635740 | JADMJW000000000 | MAG_51 | *Phenylobacterium* sp. |
| SAMN16635741 | JADMJX000000000 | MAG_54 | *Rhizobacter* sp. |
| SAMN16635742 | JADMJY000000000 | MAG_55 | *Dechloromonas* sp. |
| SAMN16635743 | JADMJZ000000000 | MAG_60 | Nitrospirae bacterium |
| SAMN16635744 | JADMKA000000000 | MAG_67 | *Vitreoscilla* sp. |
| SAMN16635745 | JADMKB000000000 | MAG_7 | *Hydrogenophaga* sp. |
| SAMN16635746 | JADMKC000000000 | MAG_77 | *Thiobacillus* sp. |
| SAMN16635747 | JADMKD000000000 | MAG_80 | *Microbacterium* sp. |
| SAMN16635748 | JADMKE000000000 | MAG_89 | *Candidatus* Sericytochromatia bacterium |
| SAMN16635749 | JADMKF000000000 | MAG_9 | *Aquabacterium* sp. |
| SAMN16635750 | JADMKG000000000 | MAG_92 | *Thiobacillus* sp. |
| SAMN16635751 | JADMKH000000000 | MAG_95 | *Thiobacillus* sp. |
